# An Automated Method for the Assessment of Memory and Learning in Larval Zebrafish

**DOI:** 10.1101/2022.06.16.496439

**Authors:** Mark Widder, Chance Carbaugh, William van der Schalie, Yuanzhang Li, Monica Martin, Brandon Pybus, Patricia Lee, Charlotte Lanteri

## Abstract

Zebrafish have shown value in translational research for many human diseases, including neuropsychiatric disorders and cognitive dysfunction, with low cost and rapid testing facilitating drug screening and discovery. However, some endpoints, such as associative learning and long-term memory, have relied historically on manual data tracking that slows data acquisition and increases costs. We automated an associative learning/long-term memory test developed by Hinz et al.^1^ using a Noldus DanioVision system and EthoVision software to enhance assay throughput and utility for monitoring learning and memory behavior through automated tracking of location and movement of individual fish.

## Article

Zebrafish have been proposed for use in translational research and drug discovery for a wide range of human diseases, including neuropsychiatric disorders^2-7^ and cognitive dysfunction^8^, and have the benefit of being a relatively low cost and rapid in vivo screening system to down-select hit compounds for further preclinical assessments. However, for certain endpoints, such as associative learning and long-term memory, tests with automated data tracking may not be available. Here, we describe methods for increasing assay throughput through automated data analysis for the associative learning/long-term memory test developed by Hinz et al.^1^ that originally used manual scoring of zebrafish position to demonstrate learned behavior.

In the zebrafish associative learning model, zebrafish larvae (6-8 days post-fertilization) are trained to associate an adverse spatial location (dark side of a chamber) with a social reward: a clear window to view conspecific fish. Larval fish are initially transferred individually into chambers separated by opaque infrared-transparent dividers (Supplemental Figure 1). After a 5-minute acclimation period, a video projector superimposes light or dark areas (Supplemental Figure 2) on alternative sides of the test chamber for 15 minutes. This establishes the normal light/dark preference pattern for each fish; typically, the light side is preferred (Supplemental Figure 3). In the original Hinz et al.^1^ method, the position of the fish was recorded and subsequently manually checked every 10 seconds. In our improved method, both the position and movement of each fish in the light and dark areas are automatically collected with a DanioVision system using Noldus EthoVision software (Figure 1A).

**Figure 1.**
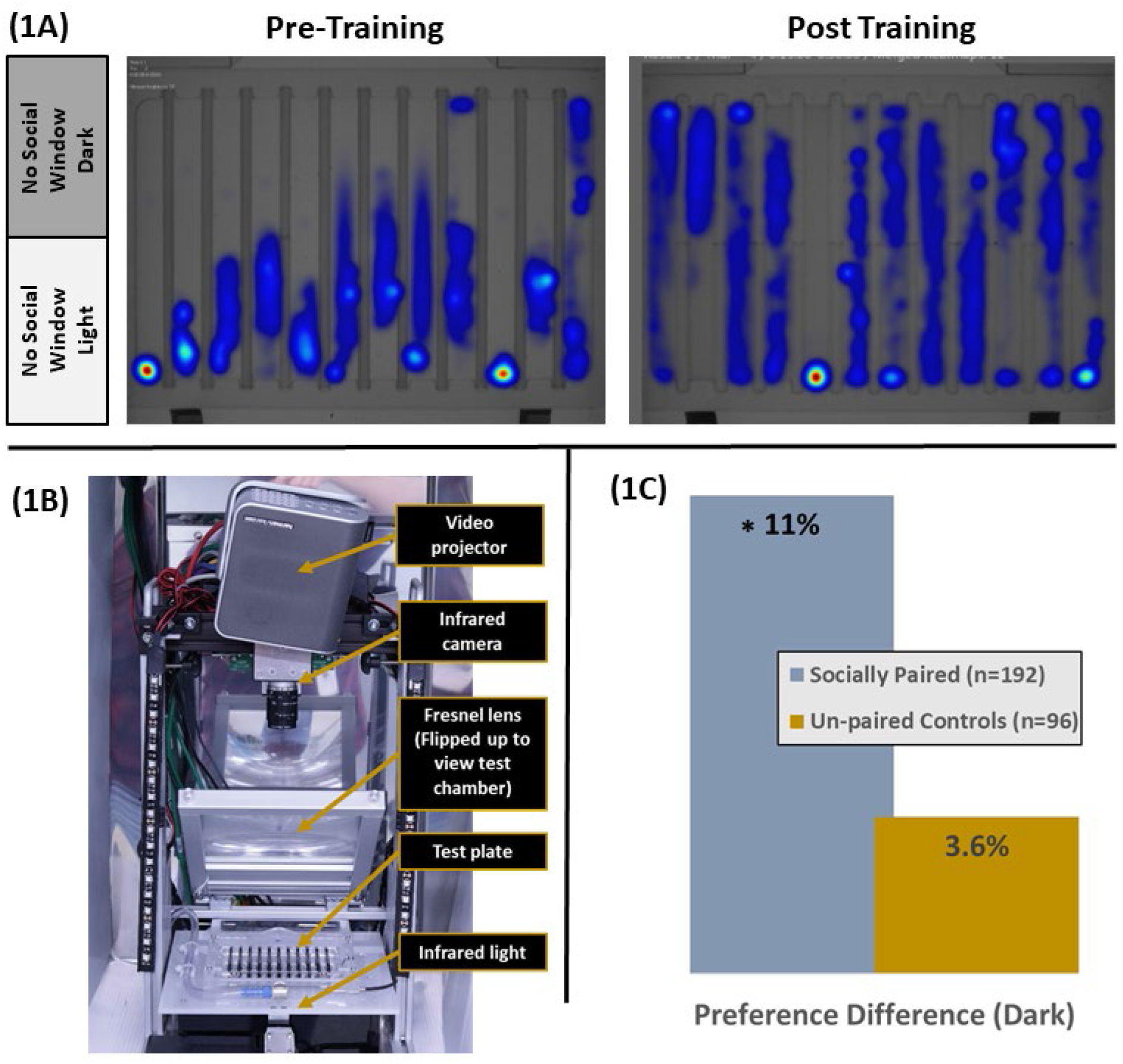
1A. Fish tend to spend most of the time in the light when other fish cannot be viewed (pre-training), but increase time in the dark when trained to associate the dark with the presence of other fish (post-training). 1B. DanioVision System used to conduct assay. 1C. Changes in amount of time spent in the dark from pre-training to post-training, showing retention of memory associating a dark environment with prior social reward. * Significance at p=0.04, t-test

Following the 30 minute pre-training assessment, the fish are trained to associate a view of conspecific fish with the dark side of the chamber. For training, fish (socially paired) are transferred into a chamber with half-clear and half-opaque infrared-transparent dividers, which is used so that fish can view other fish in the clear half of the chamber (Supplemental Figure 4). The clear side of the chamber is kept in the dark, an environment the larval zebrafish would normally avoid. Fish are trained for 3 hours, using 45 minute alternating light and dark projections. The plates are manually rotated every 45 minutes as the video projection switches to match the social window with the dark side of the chamber and avoid any chamber position bias (Supplemental Figure 5). After training, fish are transferred to individual wells of a 12-well microplate and held overnight. At 24 hours post-training, the fish are placed back into the chamber with opaque infrared-transparent dividers and post-training monitoring is then conducted for 30 minutes following the same procedures as for pre-training (Supplemental Figure 6). For comparison, un-paired control fish (control fish trained with no social window) are subject to the same procedures, but are held in chambers with opaque dividers throughout, including during the training period. Figure 1B shows how fish increase their time in the dark area of the chamber after training, even when conspecific fish cannot be viewed.

Dark preference and preference difference (pre-versus post-training) were calculated for both socially paired and unpaired control groups:

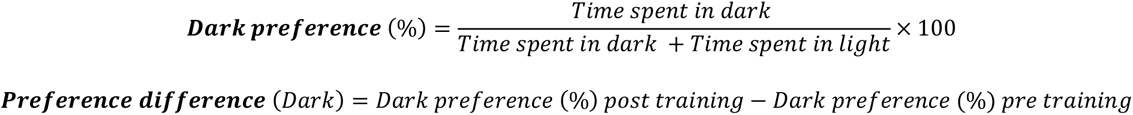

A paired t test and General Linear regression model adjusted by chamber location and training time were performed on the preference difference (dark) (Supplemental Note 1). Similar to Hinz et al.^1^, we found that 24 hours after training, socially paired fish showed a significant change in preference (11.0%, p=0.04) for the dark environment when compared to pre-training. Unpaired controls showed a 3.6% difference to the same conditions, which was not significant (Figure 1C). In addition to location preference, automated tracking allowed monitoring of movement/activity, and socially trained fish also showed significantly greater movement (p=0.03) in the dark environment after training when compared to unpaired controls (Supplemental Figure 7). This increased movement in the dark environment suggests that trained fish are searching for social cohorts in the dark environment even 24 hours after training.

In conclusion, we improved the Hinz et al.^1^ method by automated tracking of fish position and movement using the commercially available DanioVision system and EthoVision software. Additional efforts are underway to further improve the throughput and utility of the model, including the use of tablet computers to simultaneously train multiple test chambers. The methods developed here facilitate rapid assessment of memory and learning in zebrafish, enabling broader screening and discovery of interventions against cognitive dysfunction.

## Supplemental Notes

Supplemental Note 1.

The dark preference variable before and after training was paired for the same fish, and the paired t test and univariate analysis was used to examine the difference of the mean preference between control (non-trained fish) and cases (trained fish) and also to examine the data distribution and variances equivalence between cases and controls. Since other factors, such as fish location in the test chamber and fish training time (morning or afternoon) may have affected fish behavior, a General Linear regression model adjusted by location in the test chamber and training time was performed on the preference difference (dark).

Preference Difference (Dark)=intercept+ β_1*if case+β_2*location+β_3*Time+ε

The regression coefficient of β1, which showed the adjusted Preference difference (Dark) by control the location and time factors.

Socially paired fish showed a significant change in preference (11.0%, 95% CI (6.81, 15.18) p=0.04)) for the dark environment when compared to pre-training. Unpaired controls showed a non-significant change (3.6%, 95% CI (−1.94, 9.14)).

## Supplemental Figures

**Supplemental Figure 1.**
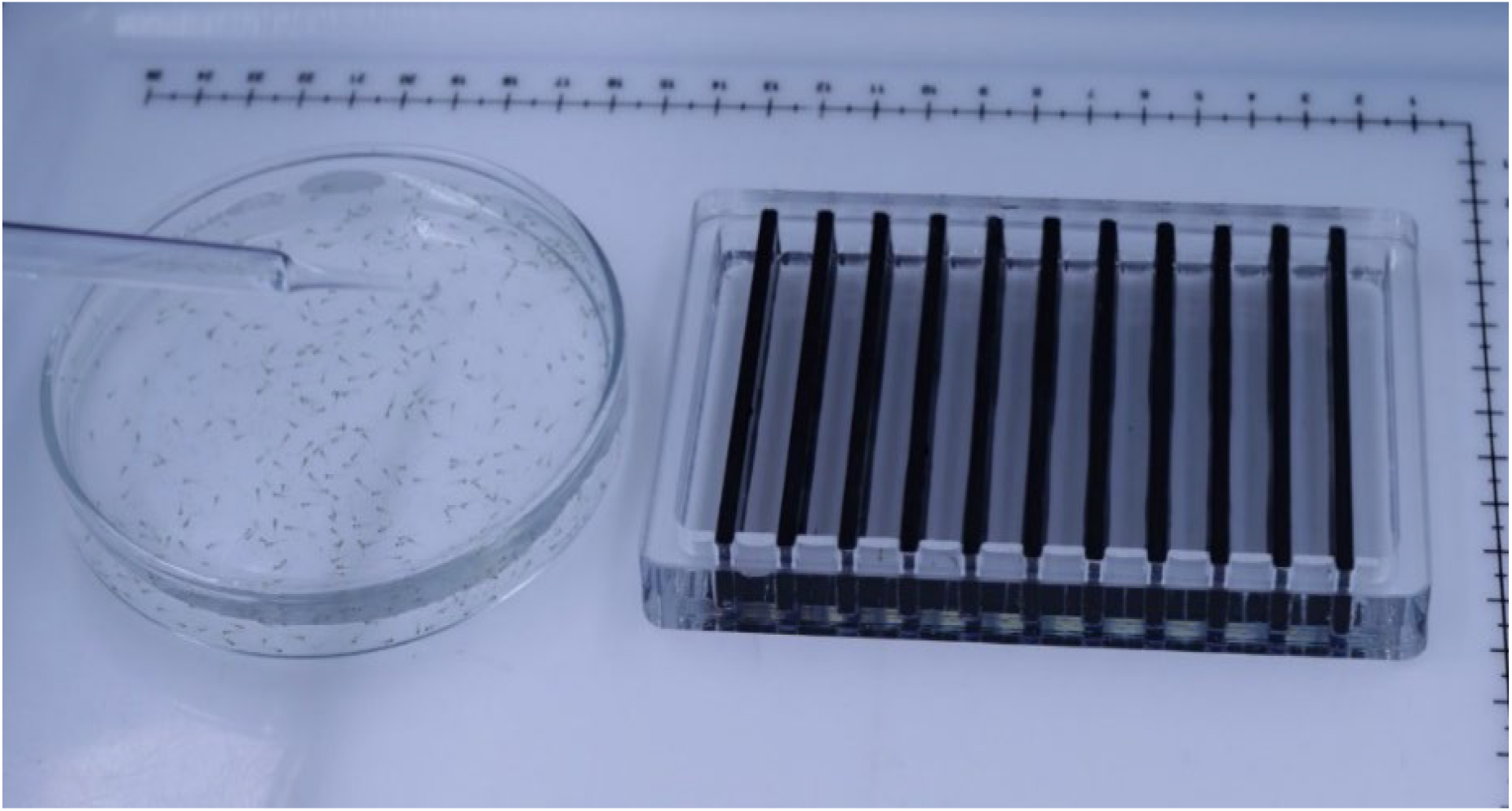
Larval fish shown next to test chamber with opaque infrared-transparent dividers. This chamber is used for pre- and post-training 30-minute

**Supplemental Figure 2.**
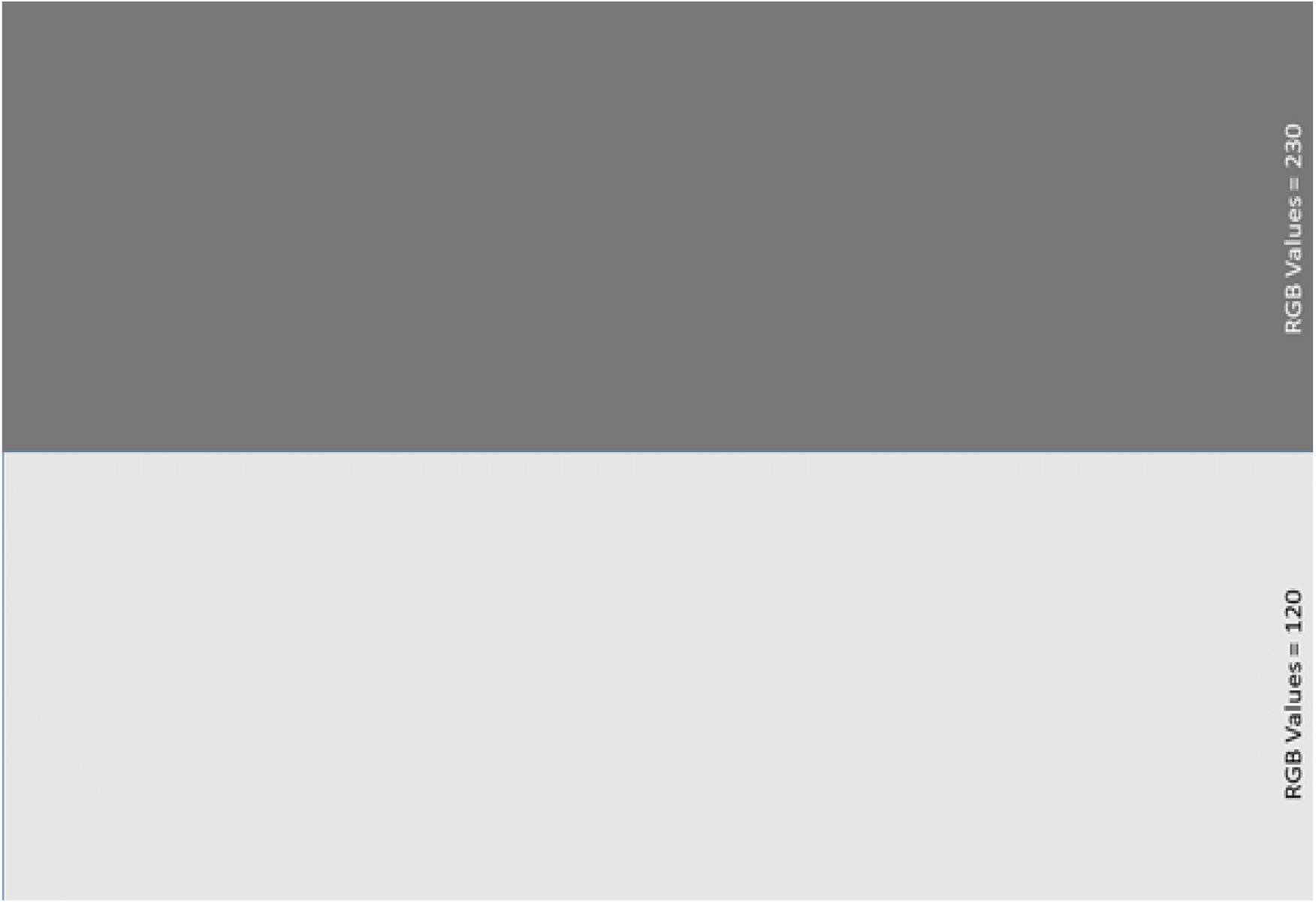
Video projection used to superimpose light and dark areas onto testing chamber.

**Supplemental Figure 3.**
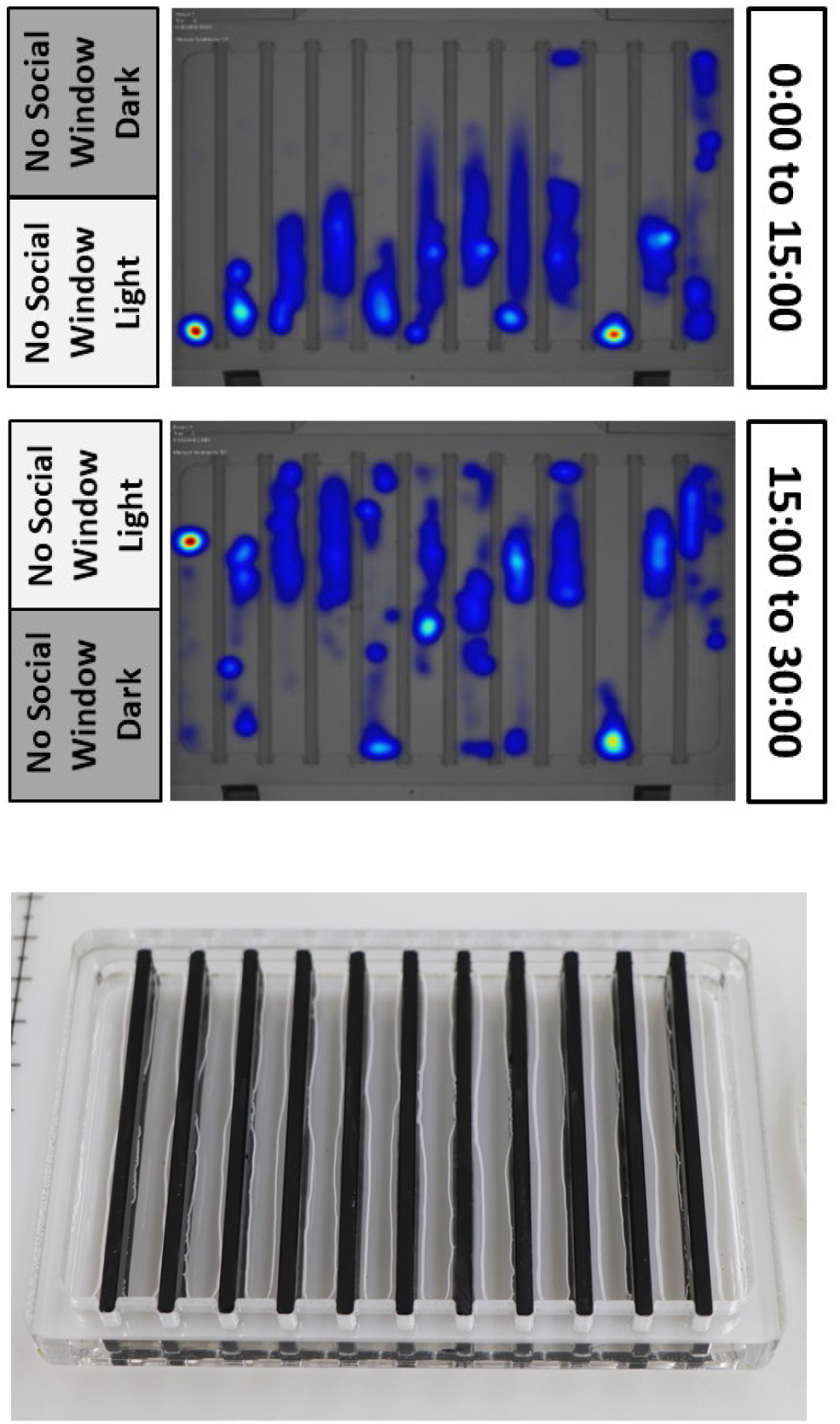
Pre-training (30 minutes). Fish are in chambers with opaque infrared-transparent dividers. With no social window, fish stay away from the dark side of the chamber.

**Supplemental Figure 4.**
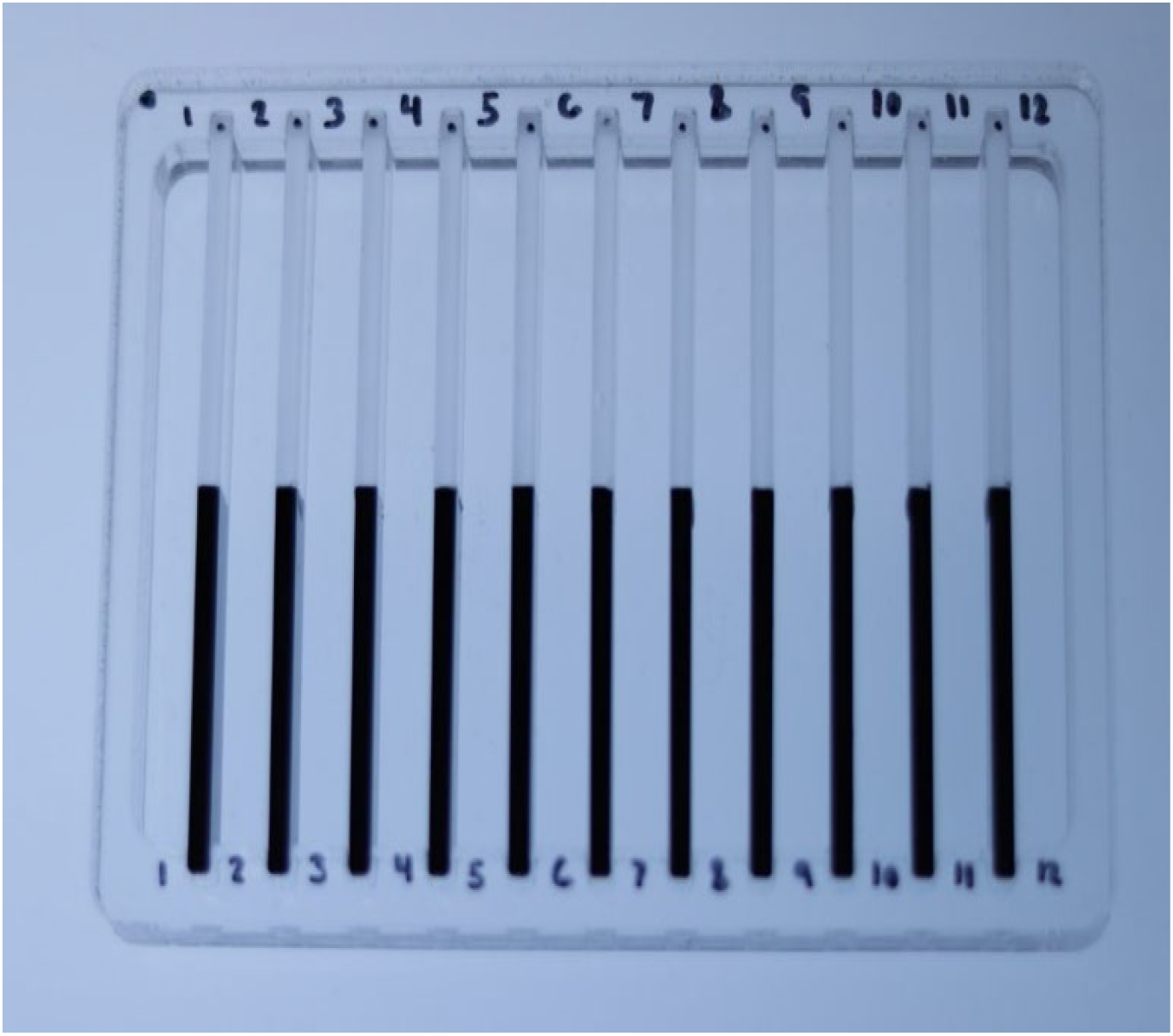
Chamber with half clear and half opaque infrared-transparent dividers used for 3 hour social reward training.

**Supplemental Figure 5.**
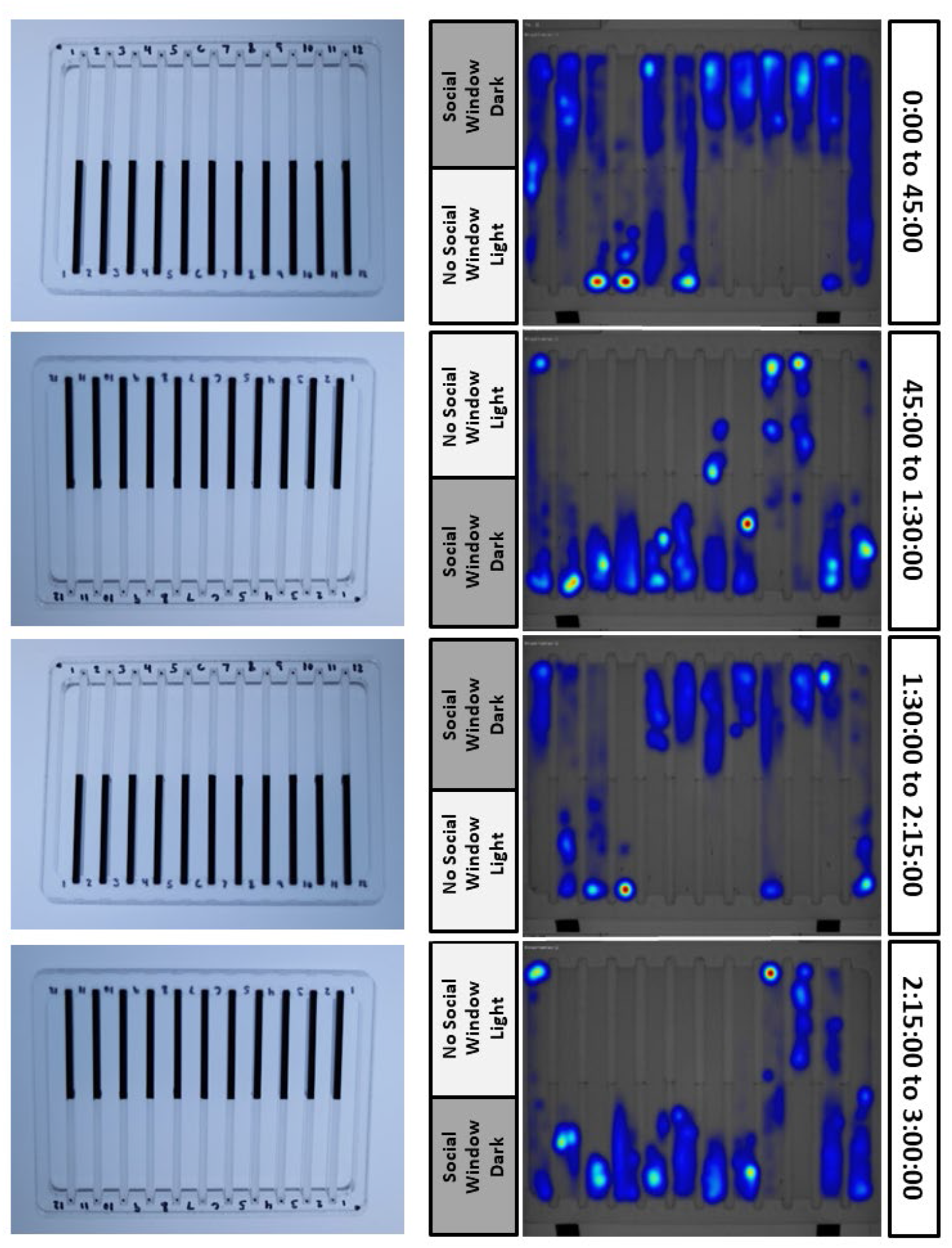
Training (3 hours). After pre-training, fish are kept in chambers with half clear sides allowing them to view other fish, but the clear side is kept in the dark. Fish learn to spend more time on the dark/clear side of the chamber.

**Supplemental Figure 6.**
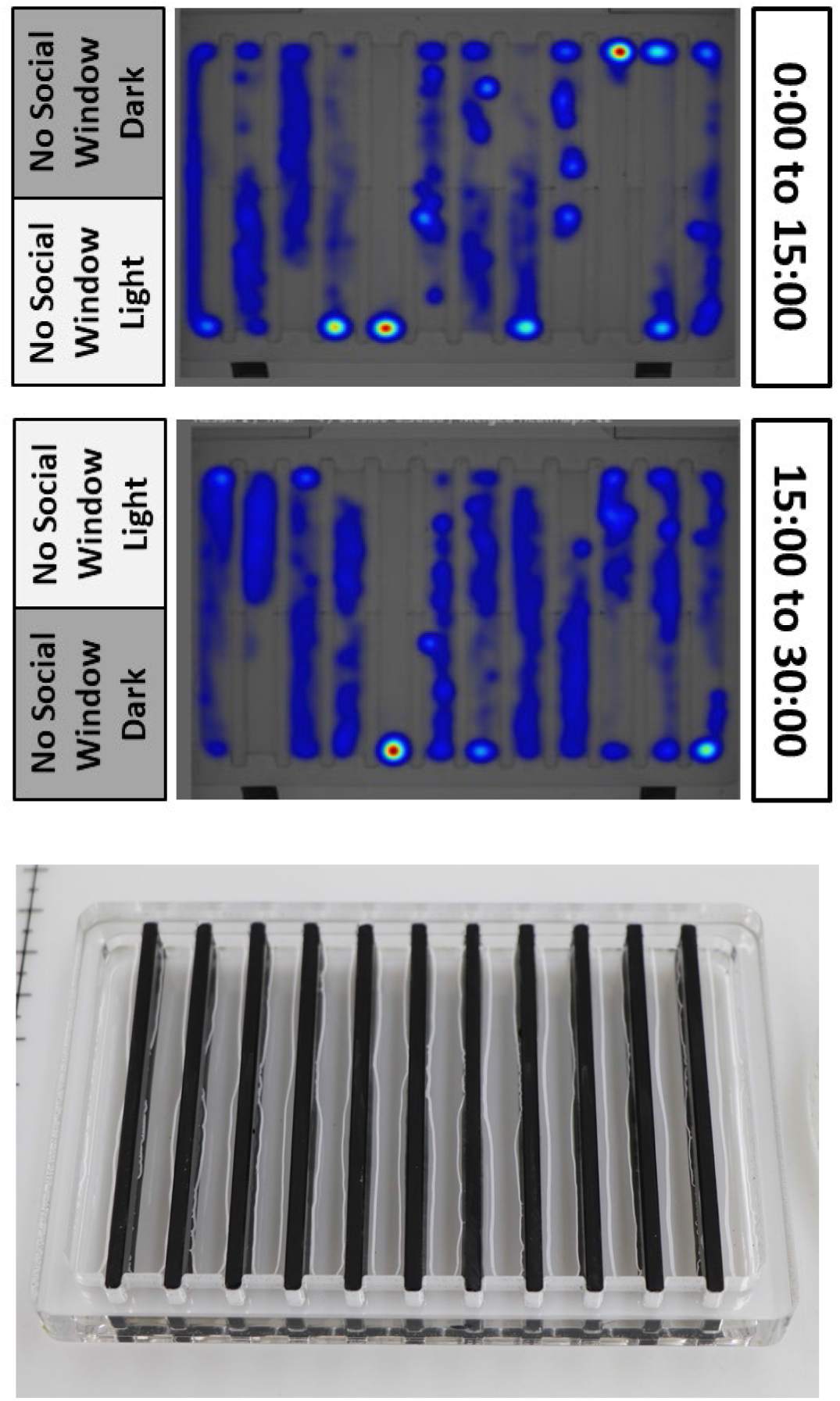
Post-training (30 minutes). To test the fish memory of learned behavior, fish are kept for 24 hours after training and the returned to the chamber with all opaque infrared-transparent dividers. The position and movement patterns are monitored as in pre-training, and the degree to which fish retained their learned behavior is determined.

**Supplemental Figure 7.**
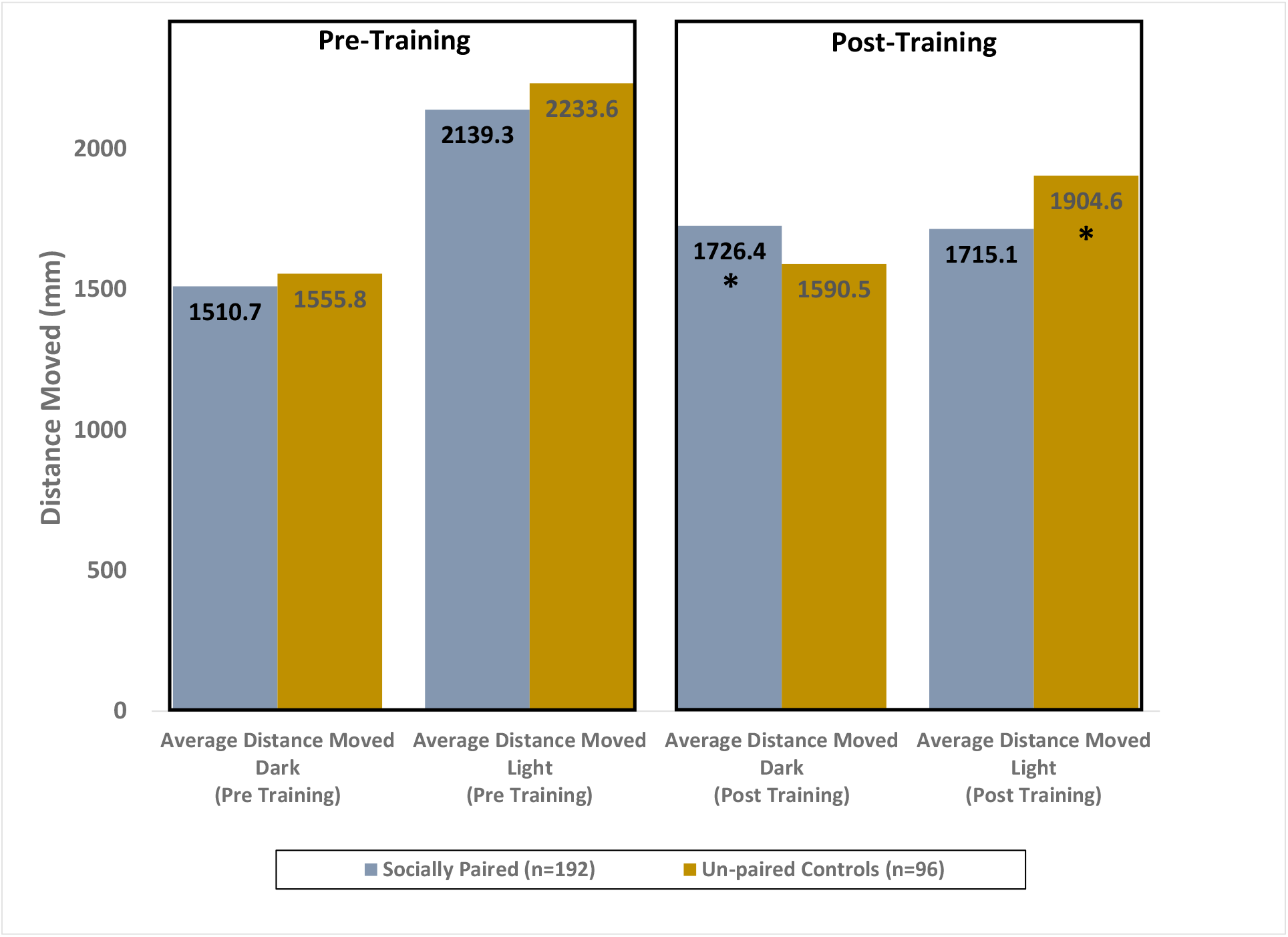
Average distance moved (mm) for socially paired and unpaired control fish during pre-training and post training on light and dark sides of the chamber. Using generalized linear regression, no difference was measured between the two groups in either the light or dark areas of the chamber during the pre-training evaluation. *During the post-training evaluation (24 hours after social training), the socially paired fish showed statistically higher movement (p=0.03) on the dark side of the chamber while unpaired fish showed statistically higher movement (p=0.04) on the light side of the chamber compared to their socially trained cohorts.

